# RNA editing of CAPS1 regulates synaptic vesicle organization, release and retrieval

**DOI:** 10.1101/178202

**Authors:** Randi J. Ulbricht, Sarah J. Sun, Claire E. DelBove, Kristina E. Kitko, Saad C. Rehman, Michelle Y. Wang, Roman M. Lazarenko, Qi Zhang, Ronald B. Emeson

## Abstract

Calcium-dependent activator protein for secretion 1 (CAPS1) facilitates the docking and priming of synaptic and dense core vesicles. A conserved hairpin structure in the CAPS1 pre-mRNA allows an post-transcriptional adenosine-to-inosine RNA editing event to alter a genomically-encoded glutamate to a glycine codon. Functional comparisons of CAPS1 protein isoforms in primary hippocampal neurons show that elevation of edited CAPS1 isoforms facilitates presynaptic vesicle clustering and turnover. Conversely, non-edited CAPS1 isoforms slow evoked release, increase spontaneous fusion, and loosen the clustering of synaptic vesicles. Therefore, CAPS1 editing promotes organization of the vesicle pool in a way that is beneficial for evoked release, while non-edited isoforms promote more lax vesicle organization that widens distribution, attenuates evoked release and eases the control of spontaneous fusion. Overall, RNA editing of CAPS1 is a mechanism to fine tune neurotransmitter release.

**IMPACT STATEMENT:** Post-transcriptional RNA editing of CAPS1 is a mechanism to regulate neurotransmitter release from synaptic vesicles.

## INTRODUCTION

The conversion of adenosine-to-inosine (A-to-I) by RNA editing is a post-transcriptional modification in which genomically-encoded adenosine residues are subject to site-specific deamination in precursor and mature mRNAs, tRNAs and primary miRNA transcripts. Cellular processes that rely upon base-pairing interactions (e.g. codon recognition during translation), recognize inosine nucleosides as guanosine residues which can alter the amino acid coding potential of RNA targets from that originally specified by genomic DNA (Higuchi et al., 1993; Sommer et al., 1991). Although only a small fraction of identified editing sites predict non-synonymous codon changes in the open reading frames of mRNAs, these recoding events often occur in transcripts encoding proteins critical for nervous system activity including ligand- and voltage-gated ion channels, G-protein coupled receptors and components of the synaptic release machinery (Hood and Emeson, 2012; Hoopengardner et al., 2003; Rosenthal and Seeburg, 2012). One such re-coding site occurs within transcripts encoding calcium-dependent protein for secretion 1 (CAPS1) (Li et al., 2009), an essential gene involved in the regulated release of hormones, neurotransmitters and peptides from synaptic vesicles (SVs) and dense core vesicles (DCVs) (James and Martin, 2013).

CAPS1 was originally characterized as a brain-derived cytosolic factor required for calcium-mediated release of dense core vesicles from permeabilized adrenal pheochromocytoma (PC-12) cells (Ann et al., 1997; Walent et al., 1992). Accumulating evidence has suggested that CAPS1 acts as a priming factor, stabilizing the interaction of vesicle- and plasma membrane-associated SNARE proteins (Grishanin et al., 2004; James et al., 2010; Sugita, 2008), to facilitate formation of the trans-SNARE complex (James et al., 2010) through interaction of its multiple domains with exocytotic factors. The essential pleckstrin homology domain within CAPS1 interacts with phosphatidylinositol 4,5-bisphosphate (PIP2), a membrane phospholipid enriched in zones of the membrane that are actively undergoing release events and this CAPS1/PIP2 interaction is essential for CAPS1 function and its localization to the active zone (James et al., 2010). CAPS1 also contains a Munc homology domain that facilitates interaction with components of the SNARE complex, including syntaxin 1 (Betz, 1997), SNAP25 and VAMP2 (Daily et al., 2010), while the C2 membrane interaction domain is required for membrane-dependent dimerization of CAPS1 which contributes to its exocytotic activity (Petrie et al., 2016). The carboxyl-terminal domain (CTD) of CAPS1 is essential for its ability to facilitate vesicular release and is sufficient for interaction and co-localization with DCVs in PC-12 cells (Grishanin et al., 2002; Kabachinski et al., 2014). A-to-I editing occurs within a region of the CAPS1 RNA encoding the CTD to alter the identity of amino acid 1258 in the encoded protein (E/G site, mouse) (Li et al., 2009). Editing has recently been shown to increase CAPS1 interaction with the open conformation of syntaxin, and sole expression of the edited CAPS1 isoform in mutant mice increases secretion from DCVs (Miyake et al., 2016).

Here, we investigate the conservation and mechanism of CAPS1 editing, identifying a specific base paired RNA structure that is required for the site-selective editing of CAPS1 pre-mRNA. The role for CAPS1 editing in the nervous system is examined by manipulating the balance of edited and non-edited isoforms in primary cultures of rat hippocampal neurons. We find that slight alterations in the relative ratio of CAPS1 isoforms derived from edited and non-edited transcripts affects synaptic morphology, as well as presynaptic SV distribution, motility, release mode and retrieval. Consistent with previous observations, our results suggest that the edited isoform of CAPS1 increases release activity (Miyake et al., 2016). Specifically, edited isoforms of CAPS1 stabilize a population of vesicles that are tightly localized and undergo high-efficiency, activity-dependent release. However, an increase in the relative expression of the non-edited CAPS1 isoform widens SV distribution, slows evoked release, increases spontaneous release, and promotes fast and inefficient fusion events. Ultimately, these studies demonstrate a role for RNA editing in the fine-tuning of synaptic transmission and presynaptic plasticity.

## RESULTS

### Characterization of CAPS1 RNA editing

Transcripts encoding CAPS1 were initially identified as a target of A-to-I RNA editing by comparison of genomic sequences to human and mouse brain-derived cDNAs (Li et al., 2009). Consistent with recent studies that determined CAPS1 undergoes A-to-I editing in human and mouse (Miyake et al., 2016), we detect CAPS1 editing in human and mouse brain, but also sequence comparisons of RT-PCR amplicons generated from whole brain, heads or whole body from other vertebrate (rat, zebrafish) and invertebrate (*C. elegans*, *Drosophila melanogaster*) species revealed that editing of CAPS1 at this site is conserved in vertebrate species, but not in the invertebrates examined (Figure 1A-B). Next, CAPS1 RNA editing levels were quantified by a highly sensitive and accurate detection method (Hood et al., 2014) from selected mouse brain regions (hippocampus, cerebellum and frontal cortex), showing that the editing of CAPS1 RNAs is greatest in the cortex (23.5 ± 0.81%) and hippocampus (17.65 ± 0.12%), but substantially lower in the cerebellum (9.22 ± 0.21%; Figure 1C). CAPS1 editing in endocrine tissues indicates very little editing in the ovary and testes (2.06 ± 0.92% and 0.57 ± 0.05%, respectively), whereas moderate levels of editing are observed in adipose tissue, pancreas, pituitary, and thymus (10.38 ± 2.88%, 12.32 ± 4.24%, 8.73 ± 1.03%, and 8.53 ± 1.00%). Higher levels of CAPS1 editing are seen in the heart, adrenal gland and mammary tissue (27.42 ± 1.63%, 17.73 ± 2.08%, and 17.47 ± 8.00%; Figure 1C). The observed levels of CAPS1 editing in mouse largely agree with the analyses of Miyake et al. (2016) in adrenal, pancreas, cerebellum, and cortex, however, our results indicate more than twice the extent of editing for CAPS1 RNAs in the heart, a discrepancy that could result from differences in mouse strain.

**Figure 1.**
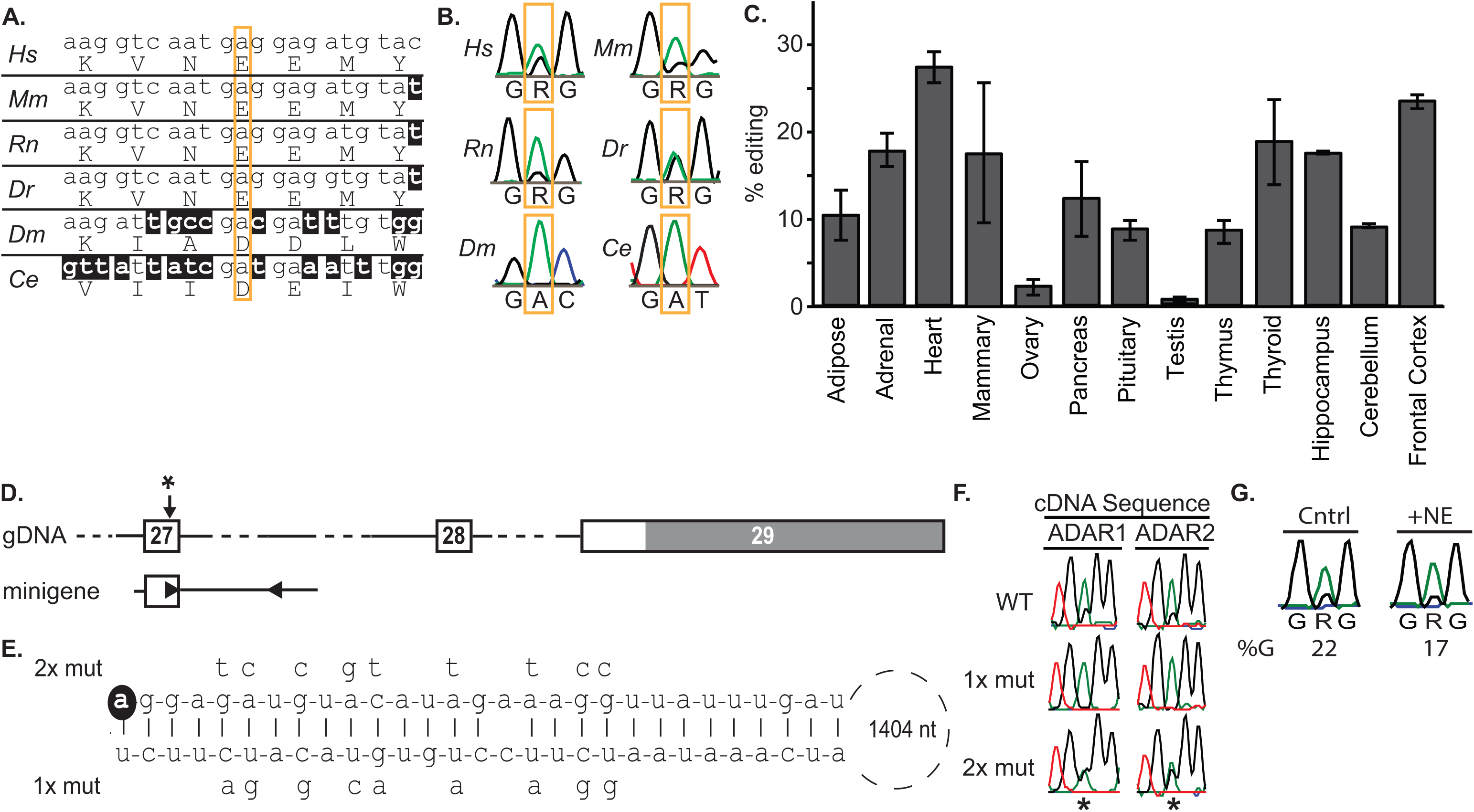
CAPS1 RNA editing in vertebrates. **A.** Alignment of CAPS1 genomic DNA and amino acid sequences. The putative editing site is indicated by a gold box from human (*Homo sapiens*, *Hs*), mouse (*Mus musculus*, *Mm*), rat (*Rattus norvegicus*, *Rn*), fruit fly (*Drosophila melanogaster, Dm*), zebrafish (*Danio rerio*, *Dr*), and nematode (*Caenorhabditis elegans, Ce*). Nucleotides not conserved with the human genomic sequence are indicated with inverse lettering. **B.** Representative electropherogram traces from Sanger sequencing of cDNAs generated from RNA isolated from whole brain (human, mouse and rat), head (zebrafish and fruit fly) or whole body (nematode). The putative editing site is indicated with a gold box; R, purine. At least three technical replicates were completed for each sample. **C.** Quantitative analysis of CAPS1 RNA editing levels in mouse tissues. The percentage of edited CAPS1 transcripts (mean ± SEM) was quantified by high-throughput sequencing analysis in RNA isolated from dissected endocrine tissues (n = 3 mice) and brain regions (n = 4 mice). **D.** Structure of the mouse CAPS1 gene from exons 27-29. The position of the editing site is indicated (*) and the 3’-UTR region in exon 29 is shaded gray. The minigene derived from mouse CAPS1 genomic DNA is shown with the locations of inverted repeats denoted (▸ and ◂). **E.** Predicted dsRNA structures formed from intramolecular base-pairing between imperfect, inverted repeats in exon 27 and intron 27 are presented where the editing site is shown with inverse lettering and the size of the loop between the inverted repeats is indicated in nucleotides (nt). Mutations to disrupt the predicted duplex structure (1X mut) are indicated below the duplex, while compensatory mutations to restore base-pairing interactions (2x mut) are indicated above. **F.** Representative electropherogram traces are shown from Sanger sequencing of RT-PCR amplicons generated from transcripts encoded by wild-type CAPS1 minigene, and mutant minigenes (1x mut and 2x mut) upon co-expression with ADAR1 or ADAR2. The position of the editing site is indicated (*); n = 3 independent transfections. **G.** Representative electropherogram traces from Sanger sequencing of cDNAs generated from RNA isolated from control (Cntrl) or primary hippocampal cultures transfected with the non-edited CAPS1 cDNA (+NE). The putative editing site is indicated; R, purine. The percentage of editing (% G) was calculated as the percentage of area under the G peak (—) divided by the total area under both the A (—) + G (—) peaks.

A-to-I RNA editing requires the presence of an extended RNA duplex for recognition by double-stranded RNA-specific adenosine deaminases (ADARs) 1 and 2 (Bass, 2002). To identify the *cis*-active sequences required for site-selective A-to-I conversion in CAPS1 transcripts, we used an RNA folding algorithm, mfold, to identify potential duplex structures containing the region of the E/G editing site (Zuker, 2003). A small, yet stable hairpin structure is predicted to occur between *cis-*elements in CAPS1 exon 27 and ~1.5kb downstream in intron 27, similar to the dsRNA region described by Miyake and colleagues (Miyake et al., 2016). To assess whether this predicted RNA duplex is required for CAPS1 editing, a genomic region containing exon 27 and ~1.7 KB of the following intron was cloned into a mammalian expression vector, and then co-transfected into HEK293T cells with cDNAs encoding the editing enzymes ADAR1 or ADAR2. RNA editing was assayed by reverse-transcription polymerase chain reaction (RT-PCR) amplification of the CAPS1 substrate, followed by sequencing (Figure 1D). The extent of editing for the wild-type minigene was ~20% for both ADAR1 and ADAR2 (Figure 1F). Mutations to the intronic region that disrupt the duplex structure (1x mut; Figure 1E) eliminate the ability of the minigene RNA to be edited by both ADARs (Figure 1F). Compensatory mutations to the exonic region of the minigene that restore the hairpin structure (2x mut; Figure 1E) also restore the ability of ADAR1 and ADAR2 to edit the minigene (Figure 1F), confirming that the identified inverted repeat elements in the CAPS1 pre-mRNA base pair to form an RNA structure necessary for ADAR-dependent deamination.

### Editing-dependent changes in CAPS1 synaptic localization

We chose primary cultures of rat hippocampal neurons as a model system to study the consequences of CAPS1 RNA editing in the nervous system. CAPS1 is robustly expressed in the dentate gyrus, CA1, and CA3 regions of the hippocampus (Lein et al., 2007). Eighteen percent of mouse hippocampal CAPS1 transcripts were edited *in vivo* (Figure 1C), in good agreement with the 22% editing observed in rat primary hippocampal cultures (Figure 1G). To manipulate levels of CAPS1 isoforms, we generated mutant CAPS1-GFP cDNA clones with a point mutation to mimic the effects of A-to-I editing at the affected codon [non-edited (GAG), edited (GGG)]. Due to the high level of endogenous CAPS1 expression, introduction of an exogenous non-edited CAPS1 isoform to cultured neurons only decreased the total level of CAPS1 editing from 22% to 17% (Figure 1G).

Confocal micrographs indicate that CAPS1-GFP isoforms (edited and non-edited) display gross patterns of localization similar to one another and to endogenous CAPS1 (Figure 2A and B). We employed both biochemical and imaging approaches to provide a more detailed examination of the localization of CAPS1 and its editing-dependent isoforms within synapses. First, crude synaptosome preparations were fractionated by selectively solubilizing presynaptic components (Phillips et al., 2001). Western blotting of these fractions indicates that CAPS1 fractionates similar to pre-synaptic markers [synaptophysin (Syp) and Munc18] and is depleted in fractions enriched with the post-synaptic marker, PSD-95 (Figure 2C); therefore, CAPS1 localization is predominantly pre-synaptic. This data agrees well with the described synaptic role for CAPS1 (Speidel et al., 2003) and studies showing that CAPS1 depletion perturbs synaptic vesicle distribution in isolated hippocampal neurons (Shinoda et al., 2016). Next, we assessed if CAPS1 editing influenced presynaptic organization using super-resolution three-dimensional structured illumination microscopy (3D-SIM; Figure 3). Edited and non-edited isoforms of CAPS1-GFP demonstrate substantial co-localization with Syp [Pearson correlation coefficients (PCC), GFP/Syp; 0.45 ± 0.016 (non-edited) and 0.50 ± 0.019 (edited)], while Syp and PSD-95 display limited PCC (0.13 ± 0.011 and 0.11 ± 0.015 in neurons containing non-edited and edited CAPS1-GFP, respectively). These results confirm that CAPS1 proteins reside primarily in the presynaptic terminals. Intriguingly, edited CAPS1 more closely correlates with localization of the synaptic vesicle marker, Syp, than the non-edited isoform (*p* < 0.05, one-tailed *t*-test; Figure 3B).

**Figure 2.**
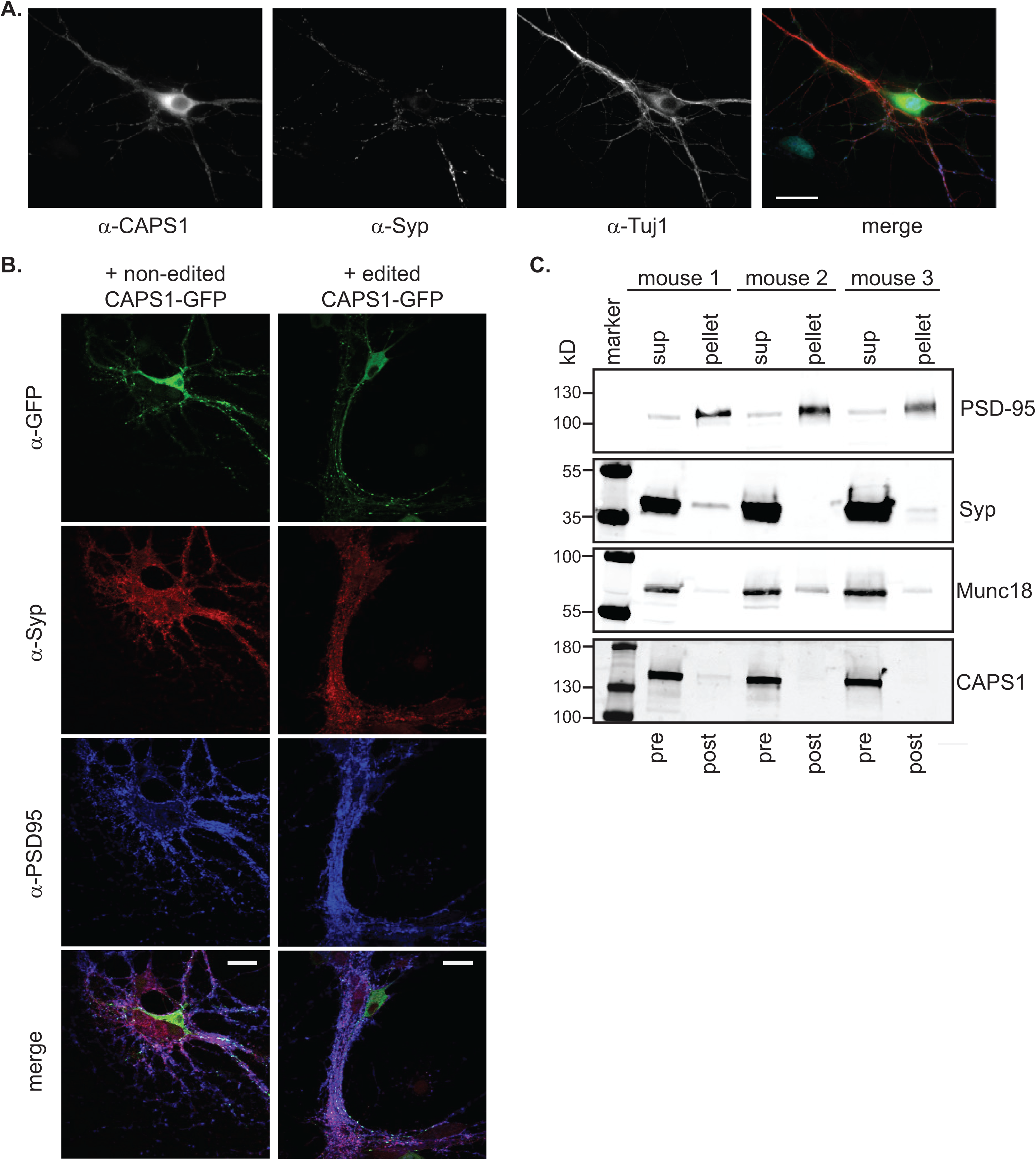
Localization of CAPS1 and synaptic proteins. **A**. Representative confocal micrographs of neurons labeled with antisera directed against endogenous CAPS1 (α-CAPS1), Synaptophysin (synapse marker, α-syp), and Tuj1 (neurite marker, α-tuj1). Scale bar = 20μm. **B**. Representative confocal micrographs of neurons transfected with non-edited (*top*) or edited (*bottom*) CAPS1-GFP isoforms and assessed for expression using antisera directed against GFP (CAPS1-GFP, α-GFP), Synaptophysin (α-Syp) and PSD-95 (α-PSD95). Scale bar = 20μm. **C.** Western blotting analysis of pre- and post-synaptic proteins in crude synaptoneurosome preparations from mouse whole brain, fractionated by extraction at pH 8.0. Extracted proteins in the supernatant (sup) represent the solubilized presynaptic fraction (pre), while post-synaptic proteins (post) remain in the pellet (pellet). Western blots were assayed using antisera directed against PSD-95, Synaptophysin (Syp), Munc18 and CAPS1 (n=3 animals).

**Figure 3.**
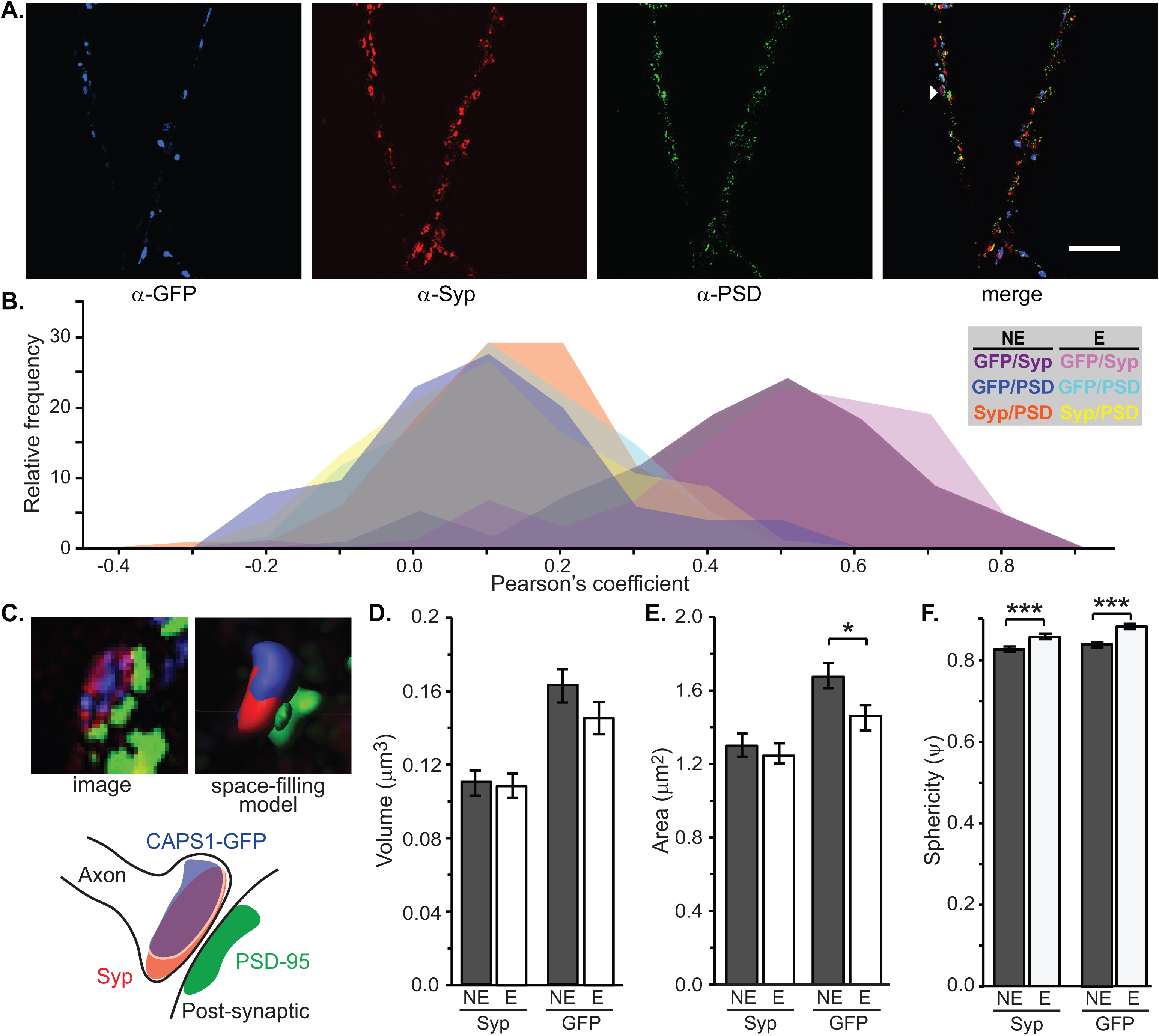
Three-dimensional structured illumination microscopy (3D-SIM) of synapses in CAPS1-GFP transfected neurons. **A.** Representative 3D-SIM micrograph of neurons transfected with CAPS1-GFP isoforms and stained with antisera directed against GFP (α-GFP), Syp (α-Syp), and PSD-95 (α-PSD). A representative synapse containing PSD-95, Syp and CAPS1-GFP is indicated (▹). Scale bar = 5μm. **B.** Ranked histogram of Pearson’s correlation coefficients from immunocytochemical co-localization analyses of Syp and PSD-95 (Syn/PSD) in the presence of non-edited (NE; purple) or edited (E; magenta) CAPS1 isoforms; CAPS1-GFP and PSD-95 (GFP/PSD) upon overexpression of NE (blue) or E (turquoise) CAPS1; and CAPS1-GFP and Syp (GFP/Syn) in the presence of NE (orange) or E (yellow) CAPS1; n = 106 (NE) and 138 (E) synapses collectively from 3 independent transfections. **C.** *Left*, representative image of synapse (indicated in panel A) containing Syp, PSD and CAPS1-GFP. *Right*, a space-filling model of an individual synapse generated from the 3D-SIM image. *Bottom*, cartoon representation of the modeled synapse showing the relative positions of synaptic components. **D-F**. Volume (D.), area (E.) and sphericity (F.) measurement of Syp- and GFP-containing loci identified as synapses in neurons transfected with non-edited (NE) and edited (E) isoforms of CAPS1-GFP; n = 208 and 202 synapses for NE or E CAPS1-GFP isoforms, respectively (from a total of 3 independent transfections); mean ± SEM; Two-tailed F-tests *p* > 0.1. Two-tailed T-tests, assuming equal variance. \**p* ≤ 0.05, \*\*\**p* ≤ 0.001. Summary data provided in Figure 3 – Figure Supplement.

Previous reports have suggested that synaptic morphology is affected by the presence of CAPS1, which primarily affects the number of SVs in the axon terminal, yet these morphological changes did not affect the area or volume of the terminal itself (Shinoda et al., 2016). Taking advantage of 3D-SIM to visualize sub-synaptic structures within 3D space, we generated space-filling models of the pre-synaptic vesicle pool structure (Syp), post-synaptic density (PSD-95) and CAPS1 localization (GFP) (Figure 3C). Focusing upon CAPS1-GFP positive synapses, we determined that the volume and area of the SV pool was unchanged by the editing status of the CAPS1 isoform expressed (Figure 3D-E). However, the Syp sphericity (ψ), a 3-dimensional measure of vesicle pool compactness (Wadell, 1935), is significantly greater in synapses expressing edited CAPS1-GFP than those expressing the non-edited (ψ_edited_ = 0.842 ± 0.006, ψ_non-edited_ = 0.812 ± 0.006; Figure 3E-F). Correspondingly, the localization of GFP is more compact with edited versus non-edited isoforms of CAPS1 (ψ_edited_ = 0.868 ± 0.005, ψ_non-edited_ = 0.830 ± 0.005; Figure 3E) and the area of edited CAPS1-GFP particles (1.46 ± 0.065 μm^2^) is significantly less than non-edited CAPS1-GFP (1.67 ± 0.067 μm^2^), yet the volume is unchanged (Figure 3D-E). These data, combined with PCC analysis, suggest that A-to-I editing increases CAPS1 co-localization with SVs and constrains SVs to a more clustered presynaptic distribution.

### Cellular impact of editing-dependent CAPS1 isoform expression

As both non-edited and edited CAPS1 isoforms co-exist in hippocampal neurons, we manipulated their relative ratios by overexpression to study the physiological consequences of CAPS1 RNA editing. Primary hippocampal cultures were transiently transfected with edited or non-edited CAPS1-GFP cDNA constructs. We probed SV release using the red styryl dye, FM4-64 (Chi et al., 2001), that is spectrally separable from CAPS1-GFP fluorescence. Transfected hippocampal neurons were loaded with FM4-64 in 90 mM K^+^, then washed. Dye uptake into presynaptic terminals expressing edited CAPS1-GFP was similar to those transfected with GFP alone or untransfected neurons in the same field of view. However, expression of non-edited CAPS1-GFP reduced the FM4-64 loading and increased the size of the synaptic bouton (Figure 4A-B). Evoked SV release was then measured by FM4-64 de-staining upon depolarization using elevated extracellular potassium. Terminals expressing non-edited CAPS1-GFP exhibited a significantly slower FM4-64 release rate than untransfected neurons, neurons expressing GFP or edited CAPS1-GFP, but there was no difference in FM4-64 loss. (Figure 4C). Therefore, non-edited CAPS1 RNA editing reduces the release probability or rate of release for recycling SVs (i.e. SVs loaded with FM4-64), but does not affect the size of the total releasable pool of SVs. Interestingly, release of styryl dyes from hippocampal neurons where CAPS1 expression has been depleted is slower than wild-type neurons (Eckenstaler et al., 2016), thus it seems that non-edited CAPS1 antagonizes CAPS1 intrinsic function for both exo- and endocytosis.

**Figure 4.**
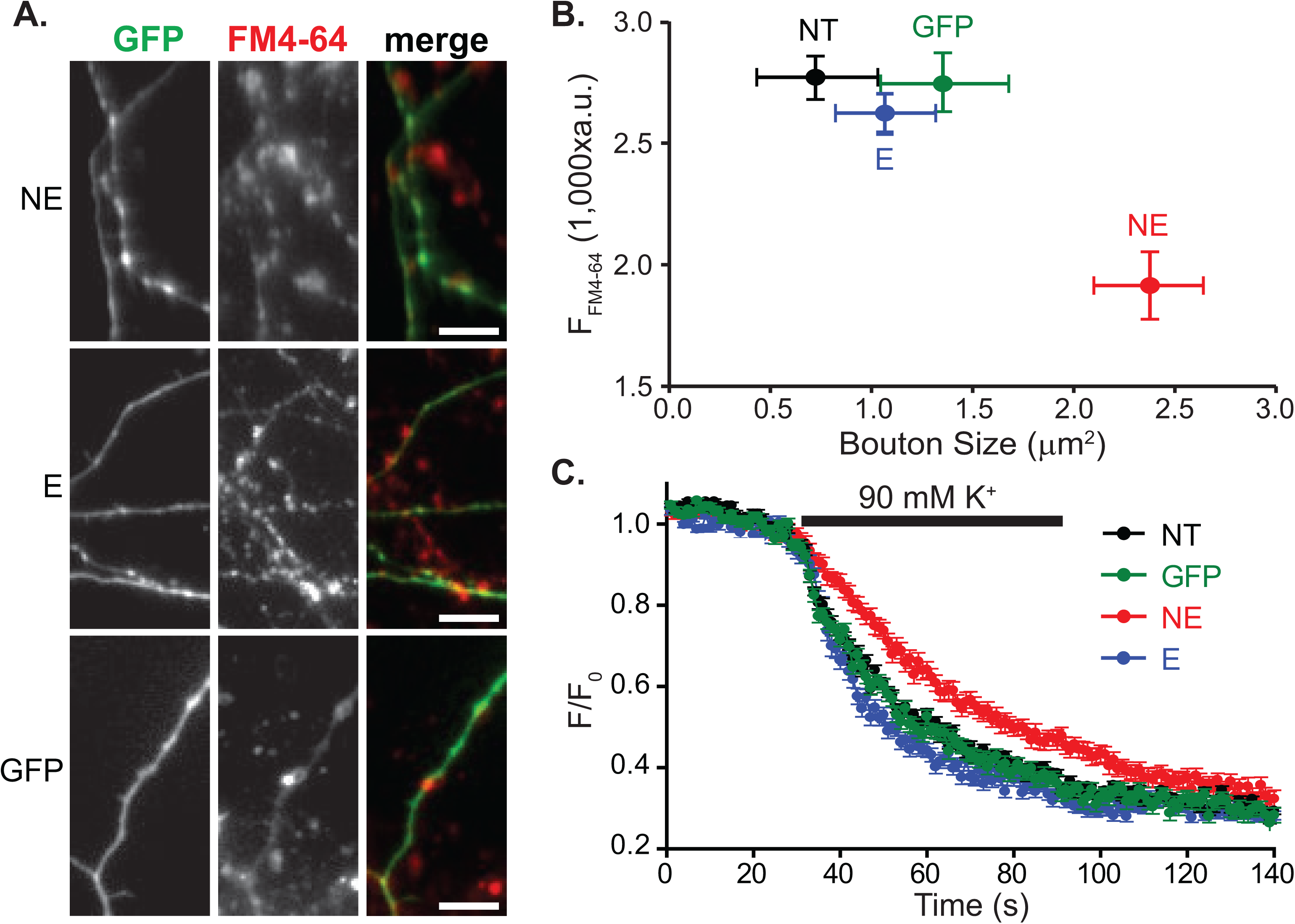
FM-dye loading and unloading of CAPS1-GFP transfected neurons. **A**. Representative images of neurons transfected with non-edited CAPS1-GFP (NE, top), edited CAPS1-GFP (E, middle) or GFP alone (bottom) and loaded with FM4-64. Scale bar = 10μm **B.** Quantification of FM4-64 dye uptake into transfected and control neurons (mean ± SEM); n=9 FOVs (fields of view) for GFP, E and NE; n=27 FOVs for NT. **C.** FM-dye (FM4-64) release. Relative fluorescence (F/F_0_ where F_0_ is the average of intensity during the baseline period before 90 mM KCl addition) from neurons during and after stimulation with 90 mM KCl solution. Neurons were either non-transfected (NT, •) or transfected with GFP (•), non-edited CAPS1-GFP (NE, •) or edited CAPS1-GFP (E, •); n = 9. Statistical analysis is provided in Figure 4 – Figure Supplement.

To further examine CAPS1 mRNA editing effects on the exocytosis and/or endocytosis of SVs, SV release and retrieval was continuously monitored using pHTomato (a pH-sensitive mCherry mutant) fused to the luminal domain of Synaptophysin (Syp), SypHTm. Exocytosis results in a luminal pH increase and thus an increase of SypHTm fluorescence, whereas vesicle re-acidification after endocytosis quenches SypHTm (Li and Tsien, 2012). In each imaging experiment, application of 50 mM NH_4_Cl elicited fluorescence by neutralizing all intracellular acidic compartments (including non-releasable vesicles) and subsequent perfusion of acidic Tyrode’s solution (pH 5.5) quenches all SypHTm, including that on the cell surface (Li and Tsien, 2012), allowing estimation of the fraction of surface and vesicular SypHTm (Figure 5A). In axon terminals expressing non-edited CAPS1-GFP, there was significantly more SypHTm fluorescence observed at rest [relative fluorescence in 4 mM KCl, pH 7.4: 0.5472 ± 0.0107 (non-edited) versus 0.5223 ± 0.0058 (control); Figure 5B-C], indicating more SV protein dwelling on the presynaptic membrane surface. By contrast, expression of edited CAPS1-GFP decreased SypHTm fluorescence at rest (0.4843 ± 0.0081), indicating that edited CAPS1 reduces surface dwelling of SV proteins (Figure 5B-C).

**Figure 5.**
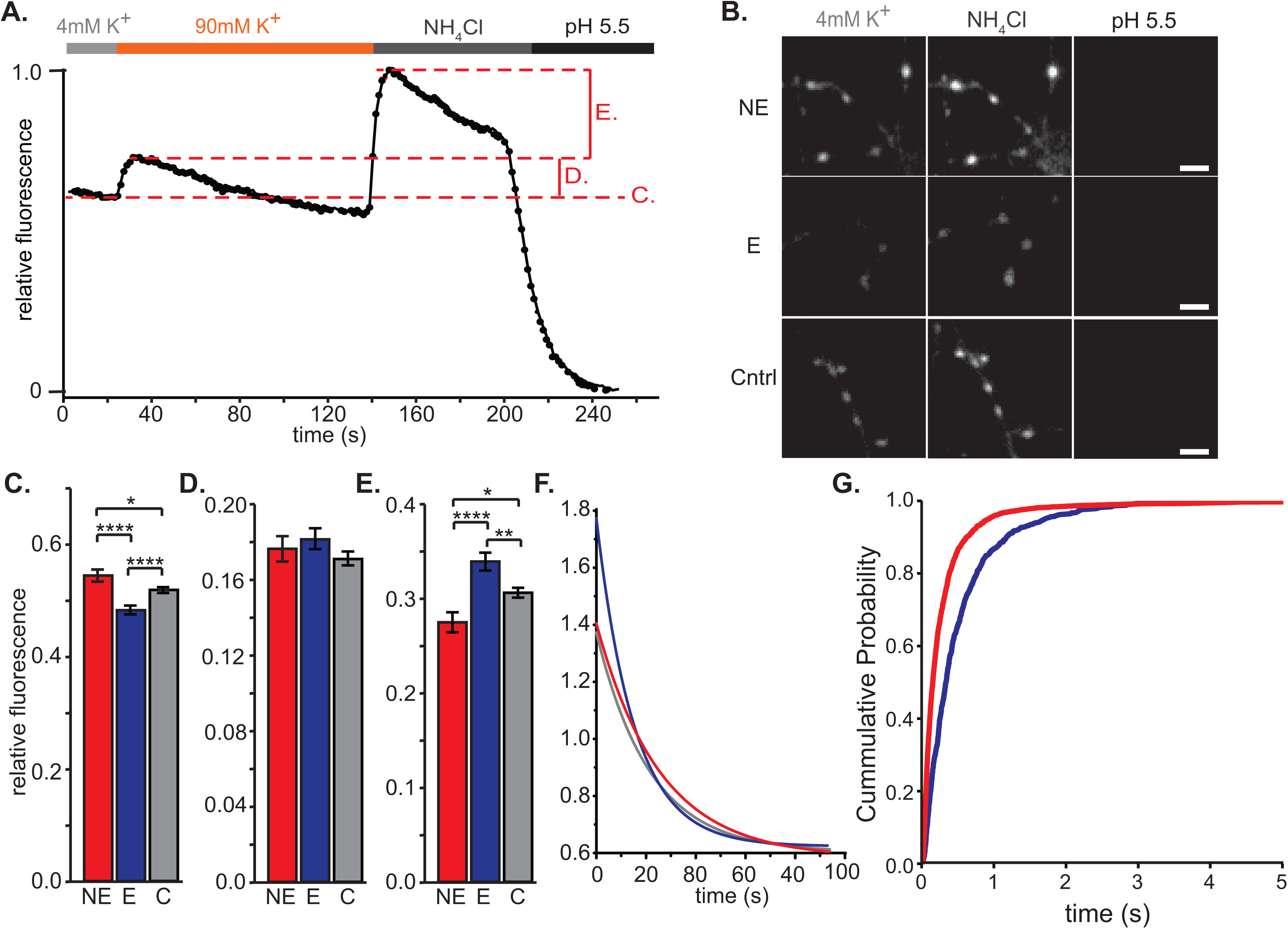
Presynaptic activity in CAPS1-GFP transfected cultured hippocampal neurons. Sample trace of relative SypHTm fluorescence from an individual synapse in a transfected hippocampal neuron under conditions of 4 mM potassium (**—**), 90 mM potassium (—), NH_4_Cl (**—**) and low pH Tyrode’s solution (pH5.5, **—**); C, D and E indicate the source of calculated data in their respective panels. **B**. Representative images of SypHTm fluorescence from neurons containing non-edited CAPS1-GFP (NE), edited CAPS1-GFP (E) or SypHTm only (Cntrl) under conditions of 4 mM potassium, NH_4_Cl and pH 5.5. **C.** Surface levels of SypHTm defined by mean relative SypHTm fluorescence at 4 mM K^+^ (± SEM); scale bar, 5 μm. **D.** Change in SypHTm fluorescence upon stimulation with 90 mM KCl (see panel A) (mean ± SEM). **E.** Resting pool of synaptic vesicles. Difference in maximum SypHTm at 90 mM K^+^ and in NH_4_Cl fluorescence (mean ± SEM). For C-E; (NE, 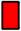) n=341 synapses from 3 independent transfections, edited CAPS1-GFP (E, 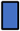) n=382 synapses from 3 independent transfections, and control (C, 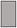) n=700 synapses from 2 independent transfections. *p ≤ 0.05, **p ≤ 0.01, ****p ≤ 0.0001. **F.** Vesicle retrieval during sustained release. Single exponential decay curves fitted to post-maximal SypHTm fluorescence under stimulatory conditions (90 mM K^+^), where the maximum signal at 90 mM K^+^ was set at 1.0. p ≤ 0.0001. **G**. mEPSC inter-event interval recorded from postsynaptic neurons which received axonal projections from GFP-outlined presynaptic cells expressing either CAPS1-GFP edited (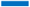) or non-edited (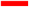); edited n=4, non-edited n=9; p ≤ 0.01. Source data and statistical analyses are provided in Figure 5 – Figure Supplement.

High-K^+^ (90mM KCl) stimulates the exocytosis of all SVs in the total releasable pool (TRP), indicated by SypHTm fluorescence (Figure 5A). In agreement with the FM4-64 result, there is no significant difference in SypHTm fluorescence with high K^+^ (control, 0.1709 ± 0.0037; non-edited, 0.1763 ± 0.0067; edited, 0.1814 ± 0.0055; Figure 5D), further suggesting that RNA editing of CAPS1 does not affect the number of SVs in the TRP. However, the fraction of SypHTm in the resting pool (i.e. SVs that fail to respond to high-K^+^ stimulation) is increased with edited CAPS1-GFP and reduced by non-edited CAPS1-GFP (control, 0.3067 ± 0.0052; non-edited, 0.2765 ± 0.0106; edited, 0.3389 ± 0.0093; Figure 5E). The resting pool is the intermediate point for endocytosed SV proteins (Rizzoli and Betz, 2005), therefore, the decrease of surface SypHTm and the increase of resting pool SypHTm by edited CAPS1-GFP indicate that RNA editing of CAPS1 promotes the endocytotic retrieval of released SVs.

Aτ the late phase of the 2-min high-K^+^ stimulation, releasable SVs are exhausted and endocytosis surges to regenerate SVs, leading to a progressive decay of SypHTm fluorescence (Figure 5A). The rate of fluorescence decay (τ) during and after high-K^+^ stimulation is faster in neurons expressing edited CAPS1-GFP [15.21 s, 95% confidence interval (CI) 14.5-15.97] versus control neurons (21.54 s, 95% CI 20.49-22.68; Figure 5F). The fluorescent decay rate is slower in neurons expressing non-edited CAPS1-GFP (25.43 s; 95% CI 24.07-26.92; Figure 5F) compared to controls. This change can be contributed to either a delay in SV retrieval rates or a more sustained SV exocytosis stimulated by the overexpression of non-edited CAPS1. The inverse situation occurred when edited CAPS1 isoforms were expressed (Figure 5C-F).

As a well-accepted measurement for spontaneous SV release, we examine the inter-event interval (the inverse of frequency) of miniature excitatory post-synaptic currents (mEPSC) with whole-cell patch clamp recording. mEPSCs are believed to be caused by spontaneous release of single synaptic vesicles from the presynaptic terminals (Lisman et al., 2007). Overexpressing edited CAPS1-GFP significantly lengthened the inter-event interval compared to non-edited isoforms (Figure 5G). These data suggest that expression of non-edited CAPS1 isoforms is reminiscent of Synaptotagmin 1 (Syt1) knockouts, where evoked release is suppressed and spontaneous release is enhanced (Sudhof, 2013). While Syt1 clamping of spontaneous release is mediated by its tailored Ca^2+^-sensitivity, edited CAPS1 can achieve similar outcomes via its reported effect of improving SV docking and priming (i.e stabilizing SVs to reduce spontaneous membrane fusion and enhance activity-evoked exocytosis) (Sudhof, 2013). This increase in spontaneous release may also contribute to the increase of SypHTm at the cell surface (Figure 5C).

While our conclusions that RNA editing of CAPS1 facilitates SV release is in good agreement with previous reports (Miyake et al., 2016), we sought to better understand how CAPS1 editing affects the coordination of SV release and recycling using quantum dot (Qdot)-enabled single vesicle imaging. Due to size exclusion, each SV can incorporate only a single Qdot through the classic endocytosis pathway (i.e. clathrin-mediated endocytosis, CME) and different modes of SV release and retrieval lead to unique Qdot fluorescence signatures (Zhang et al., 2007; Zhang et al., 2009). When neurons were stimulated extensively (90 mM K^+^ for 2 min) and presented with low (0.5 nM) or high (800 nM) concentrations of Qdots, single SVs or all SVs within the total releasable pool (TRP) were labeled, respectively (Figure 6A). In neurons expressing edited CAPS1-GFP, the high concentration of Qdots labeled ~10% more vesicles in the TRP than those expressing non-edited CAPS1-GFP [4,744.5 ± 91.7 a.u. (edited) versus 5,410.8 ± 135.5 a.u. (non-edited); Figure 6B], suggesting that edited CAPS1 promotes CME. In addition, the mean area of loaded Qdot clusters was significantly smaller for neurons transfected with edited CAPS1 (edited, 1.4174 ± 0.0839 μm^2^ versus non-edited, 1.6228 ± 0.1050 μm^2^; Figure 6B), a result consistent with the FM4-64 loading result indicating that SV distribution is affected by CAPS1 editing (Figure 4B).

**Figure 6.**
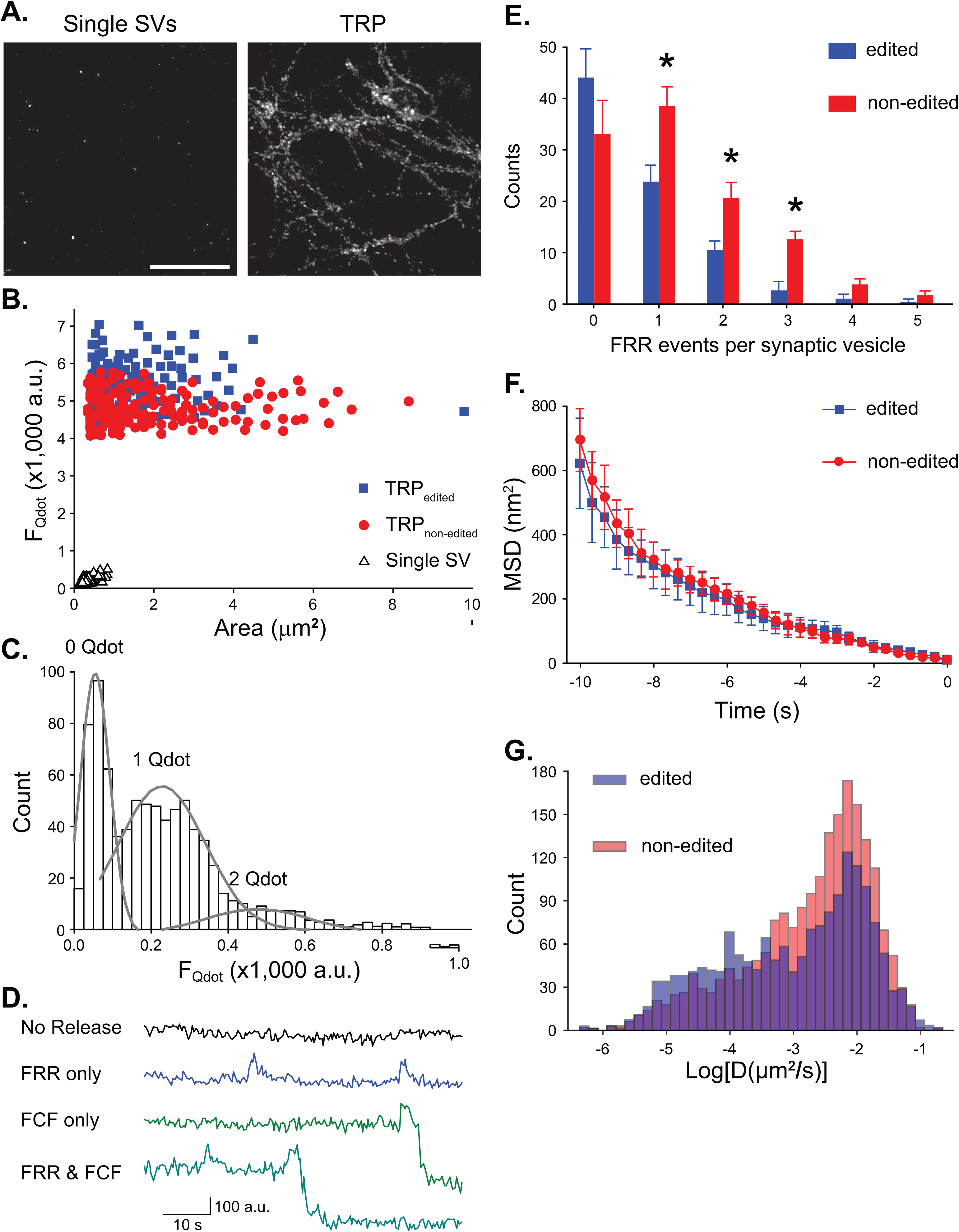
Qdot-enabled synaptic vesicle imaging at individual nerve terminals. **A.** Sample images of two different Qdot loadings resulting in randomly labeling single vesicles (*left*) or all vesicles of the total recycling pool (TRP, *right*). Scale bar, 50 μm. **B.** Distribution of Qdot photoluminescence intensity (F_Qdot_) vs. size in both loadings; the average F_Qdot_ for single Qdots is 233.1 ± 16.2 a.u. *p* ≤ 0.05, n = 9 (images from 3 neuron preparations).**C.** Quantal analysis of Qdot signal intensity at FM4-64-marked synaptic terminals after single Qdot loading. **D.** Examples of F_Qdot_ changes at FM4-64-marked synaptic terminals during 10-Hz 60-s electric field stimulation. FRR, fast release and retrieval; FCF, full-collapse fusion. **E.** Distribution of synaptic vesicles capable of conducting repeated rounds of FCF. \**p* < 0.05. Error bars represent the S.E.M. **F.** Mobility of Qdot-labeled single vesicles (measured as mean square displacement, MSD) during a 10-second period right before their first fusion (set as 0 second). **G.** Diffusion coefficient of single synaptic vesicles labeled by Qdots. *p* ≤ 0.01 n = 1350 (non-edited), n = 2147 (edited). Details on statistical analysis provided in Figure 6 - figure supplement.

At low loading concentration, Qdots randomly enter single SVs within the TRP. Photoluminescence intensity of single Qdots was independently assessed by quantal analysis (mean = 219.0 ± 34.7 a.u.; Figure 6C), matching the unitary loss of single Qdot fluorescence during exocytosis (Figure 6D). The unique patterns of Qdot signals distinguish SV turnover modes: in vsicles undergoing fast release and retrieval (FRR), like kiss-and-run, Qdots are retrieved and re-quenched, resulting in a transient and partial increase (15-30%) in their photoluminescence (Zhang et al., 2009), whereas in full-collapse fusion (FCF) a similar transient increase is immediately followed by complete loss of photoluminescence (Figure 6D), indicating release of the Qdot. During intensive stimulation (i.e. 10 Hz for 60 s), expression of non-edited CAPS1-GFP leads to individual SVs undergoing more rounds of FRR before losing Qdots to FCF (Figure 6E). In contrast, edited CAPS1 favors FCF (Figure 6E), consistent with the result (Figure 6B) indicating its promotion of CME which generally follows FCF. FCF becomes the major mode of SV release during prolonged and exhaustive stimulations, like 2-min 90-mM K^+^ perfusion or 60-s 10-Hz field stimulation, because FRR-capable SVs are in small number and used up rapidly during intensive stimulations (Gu et al., 2015; Zhang et al., 2009). Therefore, edited CAPS1 promotes efficient cargo release by FCF upon stimulation. On the other hand, FRR that is promoted by non-edited CAPS1 has a transient and narrow opening of fusion pore and thus slower and incomplete release of vesicle contents like FM dyes (Aravanis et al., 2003; Harata et al., 2006). The increase in FRR throughout the stimuli accounts for both reduced FM4-64 unloading rates (Figure 4C), and reduced τ (Figure 5F) in the presence of non-edited CAPS1, due to slowing of neurotransmitter release.

To study whether the RNA editing-related changes in TRP vesicle release and retrieval is related to the role of CAPS1 in SV docking and priming, we set to examine the mobility of TRP SVs. Taking advantage of the bright Qdot signal, we first studied the movement (mean square displacement, MSD) of SVs immediately prior to their first fusion events, a time in which docked and primed vesicles would be less mobile (Nofal et al., 2007). Surprisingly, the two different CAPS1 isoforms caused no difference in MSD up to 10 s before release (Figure 6F), suggesting that CAPS1 editing does not significantly affect SV docking and priming during the whole stimulation period.

Next, we tagged Qdots to the luminal domain of a SV protein, which allows continuous tracking of SVs, even after their release. We conjugated streptavidin-Qdots to a biotinylated antibody recognizing SV glycoprotein 2A (SV2A), a ubiquitous vesicular protein. Since there is about one copy of SV2A per SV (Mutch et al., 2011; Takamori et al., 2006), tagging SV2A should result in one Qdot per vesicle (i.e. SVs underwent exo-/endocytosis to uptake Qdots). With no exogenous stimulation, expression of edited CAPS1 resulted in reduced mobility for SVs, with mean diffusion coefficients of 7.9 ± 0.4 x 10^-3^ and 8.6 ± 0.3 x 10^-3^ μm^2^/s for edited and non-edited CAPS1 isoforms, respectively (Figure 6G). These results suggest that edited CAPS1 isoforms restrain movement of releasable/recycling vesicles in the synaptic terminal of spontaneously firing neurons. Therefore, we conclude that CAPS1 editing restricts the movement of SVs, limiting SVs to a more restricted area within the synaptic bouton (Figures 3F, 4, and 6B). Non-edited CAPS1 isoforms, on the other hand, lose such control, leading to a wider distribution of vesicles (Figures 3F, 4, and 6B). It is possible that the mobility of SVs also will affect spontaneous vesicle fusion, since more unrestricted vesicle movement could result in an increase in spontaneous interaction of individual SVs with the plasma membrane and thus increase the chance of spontaneous fusion.

## DISCUSSION

CAPS1 is required for optimal evoked exocytosis in neurons and endocrine cells, facilitating ATP-dependent priming of SVs and DCVs prior to release (Grishanin et al., 2004; James et al., 2010; Jockusch et al., 2007). A re-coding RNA editing event occurs in CAPS1 pre-mRNA transcripts and is conserved in multiple vertebrate species (Figure 1A, B). This site-specific A-to-I modification results in a glutamate-to-glycine (E/G) substitution in the carboxyl-terminal domain (CTD) of the encoded CAPS1 protein, a domain required for CAPS1 interaction with vesicles the facilitation of calcium-mediated evoked release (Grishanin et al., 2002). Our analysis of CAPS1 RNA editing in mouse tissues are largely in agreement with previous studies (Miyake et al., 2016), despite different mouse strains and different techniques employed for quantitative analysis of RNA editing profiles. Both studies identified a low level (<30%) of CAPS1 editing in mouse brain and endocrine tissues (Figure 1C), however a significant difference was observed for CAPS1 editing in the heart, where the extent of editing is 2-3 times greater that estimated by Miyake et al (2015). We empirically determined the location of *cis*-active regulatory elements required for CAPS1 editing in living cells, identifying a hairpin structure required for both ADAR1- and ADAR2-mediated A-to-I conversion in CAPS1 pre-mRNAs (Figure 1D, E, F).

Generation of genetically-modified mice that solely express the edited CAPS1 isoform (E^1252^G) recently has been shown to increase DCV release from isolated chromaffin cells and dopamine secretion from striatal synaptosomes (Miyake et al., 2016), suggesting that editing of CAPS1 enhances its role in evoked release. In the present studies, cultured hippocampal neurons were chosen as a model system where CAPS1 is highly expressed from the endogenous locus in the presynaptic compartment (Figure 2A and C), and displays an RNA editing profile similar to adult mouse hippocampus (Figure 1G), thereby providing a unique model where it is possible to manipulate the ratio of CAPS1 isoforms. Even small alterations in the ratio of edited and non-edited CAPS1 isoforms generates measurable and consistent alterations in synaptic function (Figures 4-6) and morphology (Figures 3D-F and 4B). Importantly, our work independently provides support for the conclusion that edited CAPS1 isoforms facilitate evoked DCV and SV release. As mutant mice solely expressing the edited CAPS1 isoform showed increased release from DCVs (Miyake et al., 2016), these results support the present findings where faster, more efficient evoked secretion from SVs was observed with excess edited CAPS1 (Figures 5F and 6E). Although the majority of native CAPS1 is non-edited (>70%), it is important to note that further increasing the relative expression of non-edited CAPS1 isoforms decreased evoked SV release and recycling (Figures 4C, 5F and 6E), increased spontaneous release (Figure 5C and 5G) and decreased the pool of releasable SVs (Figures 5E and 6B).

Our analyses of CAPS1 overexpression in transfected hippocampal cultures focused on synapses expressing CAPS1-GFP. Since isoform-dependent localization in the neurons was not observed (Figure 2), the editing status of CAPS1 isoforms does not appear to grossly affect CAPS1 trafficking to axon terminals. However, it is possible that the synaptic localization of non-edited CAPS1-GFP isoforms exclude the endogenous, edited CAPS1 isoforms. In this model, the decrease in overall CAPS1 function from CAPS1-GFP (non-edited) neurons may result from stoichiometric exclusion of edited CAPS1 isoforms to a limited number of interaction sites. Alternatively, since CAPS1 functions as a dimer (Petrie et al., 2016), non-edited CAPS1 may exert a negative effect on vesicle release by serving as a dominant-negative subunit in CAPS1 heterodimers. Overall, these studies suggest that tight control of CAPS1 RNA editing represents a critical parameter in determining the role of CAPS1 in presynaptic terminals.

The current studies consistently demonstrate that CAPS1 is localized to the presynaptic compartment in neurons (Figures 2C and 3B) and that RNA editing of CAPS1 transcripts results in increased clustering of the CAPS1 protein and closer confinement of SVs within the axon terminal (Figures 3D-F and 4B). Therefore, we speculate that edited CAPS1 isoforms may influence synaptic vesicle clustering at the active zone. The editing-dependent changes in SV distribution may be related to measured increases in SV distribution into the resting pool upon increases in edited CAPS1 (Figures 5A and 6B). Since edited CAPS1 does not affect the total number of vesicles in the synaptic bouton (Figure 4C and 5D) or gross distribution (Figures 2 and 3), we conclude that CAPS1 editing affects synaptic vesicle organization rather than vesicle number. Furthermore, increasing the relative expression of edited CAPS1 leads to more efficient evoked release of cargo [an overall faster release of SV content (Figure 4C) with more FCF fusion (Figure 6E)]. These results suggest a direct relationship between vesicle pool organization and evoked release and indicate that facilitation of releasable vesicles organization may improve the efficacy of vesicle release, and thus, neurotransmitter release.

In contrast to expression of edited CAPS1 isoforms, non-edited CAPS1 is more dispersed (Figure 3F), and leads to more spreading of SVs within presynaptic terminals (Figures 3F and 4B). Since SVs are significantly more mobile in the presence of non-edited CAPS1 (Figure 6G) and the total number of vesicles in the synaptic bouton remains unaffected (Figure 4C and 5D), we conclude that non-edited CAPS1 is less concentrated at the active zone and likely diminishes its ability to promoting evoked SV release. Unconfined SVs exhibiting more movement within the presynaptic terminal are more likely to feed into the pool of spontaneously releasable pool (Kavalali, 2015), which may account for the observed increases in spontaneous vesicle fusion and synaptic protein surfacing in the presence of non-edited CAPS1 (Figure 5C and G). Upon sustained stimulation, release is slowed (Figures 4C, 5F), and FRR release events are more (Figure 6G) in the presence of excess non-edited CAPS1. An increase in these incomplete release events, concomitant with a decrease in FCF, can alter the kinetics of neurotransmitter release and thus differentiate the activation of different postsynaptic receptors (Photowala et al., 2006), which is supported by our data showing slower cargo release (Figures 5F, 4C) and a decrease in the recycling pool (Figure 5B).

CAPS1 is well known for its role in vesicle docking and priming, yet Qdot tracking detected no apparent RNA editing-associated alterations in these processes (Figure 6F), suggesting that editing alters SV organization and recycling outside of its conventional role. Other studies using CAPS1 depleted hippocampal neurons also report a post-fusion role for CAPS1, noting an activity-dependent reduction in DCV fusion and BDNF release, as well as a reduction in the overall resting pool in the absence of CAPS1 (Eckenstaler et al., 2016). This study noted a similar decrease in the resting pool of SVs (Figure 5E), completeness of cargo release (more FRR events than FCC events, Figure 6E), and in vesicle recycling rates (Figure 5F) in the presence of increased non-edited CAPS1 expression. These results support the idea that the non-edited isoform of CAPS1 is less effective in the release and retrieval of SVs than its edited counterpart. The decrease in CAPS1 function may be, at least in part, due to an observed decreased in the interaction between non-edited CAPS1 and the SNARE protein, syntaxin-1A (Miyake et al., 2016).

Previous studies have defined an SV pool that is more likely to spontaneously fuse and exhibit slow evoked FM dye release (Sara et al., 2005), characteristics shared by vesicles associated with non-edited CAPS1 expression (Figure 4). A role for both ADAR and CAPS1 in regulating spontaneous release from SVs and DCVs has been described previously (Desai et al., 1988; Fujita et al., 2007; Maldonado et al., 2013). Our results show that spontaneous release is suppressed with increased CAPS1 editing (Figure 5G) and SypHTm fluorescence indicates that non-edited isoforms increase the synaptic surface distribution of vesicular proteins (Figure 5C), which may be caused by an increase in spontaneous release and/or a decrease in endocytosis. Therefore, these studies support the idea that RNA editing of CAPS1 plays a role in the organization and turnover of the spontaneous releasable SV pool.

The present studies indicate that RNA editing modifies the role of CAPS1 in SV organization, recycling and release to modulate synaptic neurotransmission. Even modest changes in the ratio of CAPS1 isoforms encoded by edited and non-edited transcripts appear to alter SV release and distribution, demonstrating the multifaceted role of CAPS1 editing in SV recycling and pool organization. Such fine-tuning by RNA editing may provide a previously unidentified layer of synaptic regulation that bridges short-term adaptability and a longer-term plasticity that is generally associated with changes in gene expression.

## MATERIALS AND METHODS

### RNA Editing Analysis

#### Identification of CAPS1 RNA editing in diverse species

Tissue from zebrafish heads, *C. elegans* (whole body), *Drosophila* heads, Sprague-Dawley rat (whole brain), as well as dissected brain regions from adult male C57BL/6J mice were harvested. Transfected and control cultured rat hippocampal neurons (Sprague-Dawley) were harvested two days after transfection. All tissues/cells were homogenized in TRIzol and RNA was isolated according to manufacturer’s instructions (Thermo Fisher). Human whole brain total RNA was purchased from Biochain. Complementary DNA (cDNA) was generated using random primers and the High Capacity cDNA Reverse Transcription kit (Thermo Fisher) and the region surrounding the CAPS1 editing site was amplified (DreamTaq DNA Polymerase, 2X Mastermix, Thermo Fisher). Parallel control reactions lacking reverse transcriptase were performed for all samples. Amplification products were purified from a 1% agarose gel by extraction of the gel (Promega SV Wizard Gel and PCR Clean-up) and subjected to Sanger sequencing.

#### Quantitative analysis of CAPS1 RNA editing in tissues

Total RNA isolated from adrenal gland, adipose tissue, heart, mammary gland, ovary, pancreas, pituitary gland, testes, thymus and thyroid of three different adult male mice (strain CD1 ICR) was obtained from Zyagen (San Diego, CA). Dissected brain regions were obtained from adult C57BL/6J male mice and total RNA was extracted as recommended by the manufacturer (TRIzol®, Thermo Fisher). cDNA was generated as described above and amplified using a two-step amplification process, as performed previously in our laboratory (Hood et al., 2014), generating amplicons suitable for direct sequencing on the Illumina miSeq platform; first-step primers (RU9-10) are presented in Appendix Table 1.

#### Analysis of RNA editing in a CAPS1 minigene

The CAPS1 minigene containing a genomic fragment was cloned from a bacterial artificial chromosome (BAC) obtained from Children’s Hospital Oakland Research Institute (CHORI) that contains 171 Kb of genomic sequence surrounding the edited region (RP23-356M17). A *Bam HI* (7.2KB) genomic fragment containing the editing site was cloned into pcDNA3.1 and further truncated subcloning a *Hind* III restriction fragment (1.7KB) into pRC-CMV3.1 (ThermoFIsher). Mutations of the minigene (mut 1x and mut 2x) were generated using an overlapping extension version of site-directed mutagenesis (Ho et al., 1989). All restriction enzymes were obtained from New England Biolabs®, Inc. The minigene was co-transfected into HEK293 cells with rat ADAR1 (p110) or ADAR2b (Singh et al., 2007) using FuGENE® HD transfection reagent (Promega), as directed by the manufacturer. After 48 hours, cells were homogenized in TRIzol® and the RNA was extracted. Total RNA was treated twice with TURBO DNase I [TURBO DNA*-free*™ Kit (Life Technologies)], cDNA was generated as described previously and the region surrounding the editing site was amplified by PCR. Amplicons were gel-purified and subjected to Sanger sequencing. ADAR1 p110 was cloned by removing the p150 translation start (*Sal* I/*Swa* I) from clone #4012994 (ATCC).

### Cloning CAPS1 cDNA and genomic fragment

The CAPS1 fragment from an Open Biosystems cDNA clone (clone ID: 30537768; GE Dharmacon) was subcloned into pBluScript (EcoRI/NotI). Edited and non-edited isoforms of the CAPS1 C-terminal domain were amplified from total mouse RNA, subcloned into pAcGFP (Clontech) then exchanged with the CTD fragment (*Sma* I/*Sac* II) of pBS-CAPS1, to generate edited and non-edited CAPS1 lacking a stop codon and 3’-UTR. A stop codon was inserted into the *Sac* II site by insertions of annealed and digested oligonucleotides. CAPS1 isoforms were cloned into pAcGFP (*EcoR* I) to generate the final CAPS1-GFP fusion constructs.

### Primary neuron culture

For all experiments, rat hippocampal cultures were prepared as described previously (Liu and Tsien, 1995), with minor modifications. Briefly, rat hippocampi (CA1-CA3) were dissected from postnatal day 0 or 1 (P0 or P1) male and female Sprague-Dawley rats and dissociated into a single-cell suspension with a 10-min incubation in Trypsin-EDTA (Life Technologies) followed by gentle trituration using three glass pipettes of sequentially smaller diameters (~ 1 mm, 0.5 mm, and 0.2 mm). Dissociated cells were recovered by centrifugation (200 x g, 5 minutes) at 4° C and resuspended in plating media composed of Minimal Essential Medium (MEM, Life Technologies) supplemented with 27 mM glucose, 2.4 mM NaHCO_3_, 1.25 μM transferrin, 2 mM L-glutamine, 0.43 μM insulin and 10%/vol fetal bovine serum (FBS, Omega). 100 μl of Matrigel (BD Biosciences, 1:50 dilution) was deposited onto round coverslips and pre-incubated at 37° C with 5% CO_2_ for ~ 2 h, then aspirated before plating 100 μl of cell suspension (200-300 cells/ mm^2^). Cells were allowed to adhere to the coverslip surfaces for 4 h before the addition of 1 ml culture media composed of MEM containing 27 mM glucose, 2.4 mM NaHCO_3_, 1.25 μM transferrin, 1.25 mM L-glutamine, 2.2 μM insulin, 2 mM Ara-C, 1 %/vol B27 supplement (Life Technologies) and 7.5 %/vol FBS. Ara-C in the culture media efficiently prevented astroglial proliferation. Experiments were performed on neurons cultured between 12 and 17 days *in vitro* (DIV), a time-frame by which synaptic transmission is well established (Liu and Tsien, 1995). Neurons were transfected after 2-10 DIV by calcium phosphate co-precipitation (Jiang and Chen, 2006). All animals were naïve and all procedures were approved by the Vanderbilt University Institutional Animal Care and Use Committee.

### Immunofluorescence Imaging

Two days after transfection, cultured neurons were fixed in 4% paraformaldehyde and washed in 1X phosphate buffered saline (PBS). Prior to antibody staining, coverslips were subjected to antigen retrieval by treatment with boiling sodium citrate buffer (10 mM Na-citrate, 0.5% Triton, pH 6) and washed with 1X PBS at room temperature. Cultures were then blocked for 1 hour in blocking buffer (5% goat serum, 1% BSA, 0.25% Triton in 1X PBS) and stained for GFP (#A10262, Thermo Fisher), synaptophysin (ab8049, Abcam) and PSD95 (#3450, Cell Signaling Technology®) diluted in blocking buffer. Cover slips were washed three times for 5 minutes each in 1X PBS and then incubated with secondary antibodies [Goat anti-mouse cy5, Donkey anti-rabbit Cy3, Goat anti-chicken Cy 2 (Jackson laboratories)] diluted 1:500 in blocking buffer for 1 hour. Each coverslip was washed three times and mounted on glass slides (Aqua Poly/Mount cat# 18606, Polysciences Inc.). Three independent hippocampal neuron cultures were transfected, stained and analyzed by microscopy for biological reproducibility. Confocal images were taken on an LSM 510 Meta inverted microscope with a 63X/1.40 Plan-Apochromat oil immersion objective lens. Ten or more 3D-SIM images were taken from each biological replicate and ≥2 synapses were analyzed within each image. Super resolution SIM images were taken on a Delta Vision OMX, with sCMOS camera at 512x512 and a 60x Plan-Apochromat N/1.42 NA oil immersion lens. Image stacks from the Delta Vision OMX 3D structured illumination microscope was used as a template for Imaris x64 7.6.0 (Bitplane; Concord, MA) to construct three-dimensional images. The Imaris software creates surfaces encompassing all points with emission intensities exceeding a manually adjusted threshold. Separate surfaces were created for each fluorescent wavelength [synaptophysin (683 nm), PSD-95 (608 nm), and GFP (528 nm)], enabling independent measurements of 3D characteristics for each component (particle) including surface area, volume and sphericity. F-tests were performed to determine that the variance between each sample set were equal (p ≥ 0.1). Two-tailed T-tests, assuming equal variance, were then performed to determine significance between each sample.

Sphericity is a measure of the degree to which a particle resembles a perfect sphere (equation 1), where the numerator is the surface area of a perfect sphere containing the same volume as the particle of interest (V_p_) and the denominator is the surface area of the particle itself (A_p_).

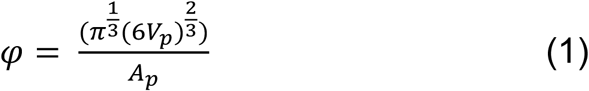

#### Live-cell Fluorescence Imaging

Two-to-four days after transfection with SypHTm and CAPS1-GFP constructs, neurons grown on glass coverslips were transferred to an inverted Nikon Ti-E microscope (40x, 1.30 NA objective, Andor iXon+ 887 EMCCD camera) equipped with a RC-26G perfusion chamber in a PH-1 platform (Warner Instruments) on an ASI 2000 motorized x-y stage. Stage and line heaters maintained the cells at 34°C (Warner Instruments). Cells were perfused at a rate of 3 ml/min with normal 4 mM K^+^ Tyrode’s saline at baseline. To stimulate cells, 90 mM K^+^ Tyrode’s saline (60 mM NaCl, 90 mM KCl, 2 mM CaCl_2_, 2 mM MgCl_2_, 10 mM HEPES, 10 mM glucose, pH 7.35) was used. NH_4_Cl solution (50 mM NH_4_Cl, 96 mM NaCl, 4 mM KCl, 2 mM CaCl_2_, 2 mM MgCl_2_, 10 mM HEPES, 10 mM glucose, pH 7.35) was used to neutralize intracellular compartments and achieve maximum fluorescence of pHTomato. Normal 4 mM K^+^ Tyrode’s adjusted to pH 5.5 was used to quench extracellular fluorescence. 4 mM K^+^ Tyrode’s was perfused for 20 seconds followed by 120 seconds of 90 mM K^+^ saline, 60 seconds of NH_4_Cl solution, and 60 seconds of pH 5.5 saline. Solution switching was controlled by pClamp 10 software (Molecular Devices) and synchronized to the beginning of image acquisition. Image acquisition was conducted automatically using Micro-manager (UCSF) where images were taken every 1.5 seconds and image analysis was performed using Image J. At least two biological replicates (neurons cultured from different rats) were conducted with ≥ 3 coverslips for each replicate. At least 29 synapses (ROIs) were independently analyzed per coverslip (Microsoft Excel). ROIs that showed no response to NH_4_Cl (maximum fluorescence at NH_4_Cl ≤ average fluorescence at 4 mM K^+^ Tyrode’s) were excluded from analysis. Curves presented in Fig 5 (panels C and E) represent moving averages incorporating all collected data (Microsoft Excel). Tau values were determined from 90mM K^+^ values (post-maximum) fitted to non-linear regression, exponential one-phase decay (GraphPad Prism 7.0).

#### FM and Qdot Imaging

FM4-64 imaging was performed on a Nikon Ti-E microscope using an iXon 897 EMCCD (Andor) with a 100X Apo VC oil-immersion objective (N.A. 1.40). Cells on coverslips were mounted in an RC-26G imaging chamber (Warner Instruments) and bottom-sealed with a 24 x 40 mm size 0 cover glass (Fisher Scientific). The chamber was fixed in a PH-1 platform (Warner Instruments) placed on the microscope stage and gravity perfusion was controlled by a VC-6 valve control system (Warner Instruments) with a constant rate of ~50 μl/sec. All perfusion lines were connected into an SHM-6 in-line solution heater (Warner Instruments). The temperatures of both the imaging chamber and the perfusion solution were maintained at 34°C by a temperature controller (TC344B, Warner Instruments). Image acquisition and synchronized perfusion were controlled via Micro-manager software. For FM dye or Quantum dot (Qdot) loading of the evoked pool of synaptic vesicles, cells were incubated with 10 μM FM4-64, 100 or 0.8 nM Qdots (Qdot 605, Life Technologies) for 2 min in high K^+^ bath solution containing 64 mM NaCl, 90 mM KCl, 2 mM MgCl_2_, 2 mM CaCl_2_, 10 mM N-2 hydroxyethyl piperazine-n-2 ethanesulphonic acid (HEPES), 10 mM glucose and 1 μM TTX, pH 7.35 (Zhang et al., 2009). After loading, cells were washed with normal bath solution containing 10 μM NBQX and 20 μM D-AP5 for at least 10 min prior to imaging. For FM4-64 and Qdot imaging, a 480 nm laser (Coherent) and a filter combination of DiC 500LX and Em 600/20 were used. All optical filters and dichroic mirrors were purchased from Chroma or Semrock. The acquisition rate was 3 Hz for Qdots and 1 Hz for all other dyes. For each dye, images were taken with the same acquisition settings including excitation light intensity, spinning disk speed, exposure time, and EM gain. All image analyses were performed in ImageJ as described previously (Gu et al., 2015). To obtain mean Qdot photoluminescence intensities in each synaptic bouton, across all fields of view, all regions of interest (ROIs) defined by retrospective FM4-64 staining of the same field of view were projected to an average image made from the first ten frames of the Qdot image stack. FM dye images were taken in 3 batches; each batch contained 3 coverslips GFP, E and NE transfections. Fluorescence intensity of every region of interest (ROI) in every FOV was averaged to obtain an average value for each FOV. Each FOV (n) was included to obtain an overall average value.

Quantal analysis for single Qdot loading was applied as described previously (Zhang et al., 2009). To analyze the behavior of vesicles labeled by single Qdots, regions of interest (ROIs) having only one Qdot were selected. Time-dependent Qdot photoluminescence changes were extracted with a 5-frame moving window. All data were exported and processed in Excel, Matlab, and SigmaPlot.

For single particle Qdot tracking, transfected neurons were pre-blocked in 1% BSA in Fluorobrite (Thermo Fisher) for 10 min before 30 min incubation with biotinylated anti-SV2a antibody (Synaptic Systems). Neurons were then incubated for 4 min with EM 605 nm Streptavidin-conjugated Q-dots (Life Technologies) in Fluorobrite containing 90mM KCl solution. Neurons were washed 6 times to remove excess surface Qdots before transfer and mounting in 4mM KCl Tyrode solution for live cell imaging. Single particle tracking was performed on an Olympus IX-81 microscope with a Nikon Intensilight lamp source and an Andor EMCCD (Exposure time = 25 ms and EM Gain = 50). 1400 frame stacks were acquired using burst acquisition. Trajectories were generated using the TrackMate function in FiJi and exported to Matlab. MSD calculations were performed as previously described (Chang et al., 2012).

### Synaptoneurosome fractionation

Crude synaptoneurosomes were prepared and fractionated as described (Phillips et al., 2001). Briefly, whole brains were removed from adult male C57BL/6J mice (n = 3) and homogenized in 10 volumes cold lysis buffer [0.32M sucrose, 4.2mM Hepes pH 7.4, 0.1mM CaCl2, 1mM MgCl2, and protease inhibitor cocktail (Sigma Aldrich)] with a Teflon-coated dounce homogenizer for 10 strokes on medium speed. Homogenates were centrifuged at 1,000g to remove cell bodies, nuclei and debris. The supernatant was centrifuged at 10,000g for 20min, and the pellet saved and washed with lysis buffer. The crude synaptoneurosome pellet was then extracted in 100mM Tris-HCl, pH 8.0 and 1% Triton X-100 for 30 min at 4°C. The supernatant was removed and saved as the presynaptic fraction, while the pellet (post-synaptic fraction) was washed with extraction buffer and solubilized in 1% SDS in 1X PBS. Equal volumes of each fraction were electrophoresed on a 4-20% Mini-PROTEAN TGX precast gel (Bio-Rad), transferred to nitrocellulose membrane (Protran; PerkinElmer), blocked (Li-Cor® Odyssey® blocking buffer; Licor Biosciences) and probed for CAPS1 (ab32787, Abcam®), Munc18-1 (#13414, Cell Signaling Technology®), PSD-95 (#3450, Cell Signaling Technology®), and Synaptophysin (ab8048, Abcam®) using secondary antisera consisting of fluorescently labeled anti-mouse (#926–32212), anti-goat (#926– 68024), and anti-rabbit (#926–32213) immunoglobulin (Licor® Biosciences) suitable for imaging on the Li-Cor® Odyssey® infrared imaging system. Images were quantified using ImageJ.

### Electrophysiology

Whole-cell voltage clamp recordings were performed on neurons from 12 - 17 DIV cultures using a Multi-Clamp 700B amplifier, digitized through a Digidata 1440A, and interfaced via pCLAMP 10 software (all from Molecular Devices). All recordings were performed at room temperature. Cells were voltage clamped at −70 mV for all experiments. Patch pipettes were pulled from borosilicate glass capillaries with resistances ranging from 3 - 6 MΩ when filled with pipette solution. The bath solution (Tyrode’s saline) contained: 150 mM NaCl, 4 mM KCl, 2 mM MgCl_2_, 2 mM CaCl_2_, 10 mM N-2 hydroxyethyl piperazine-n-2 ethanesulphonic acid (HEPES), 10 mM glucose (pH 7.35). The pipette solution contained: 120 mM cesium methanesulfonate, 8 mM CsCl, 1 mM MgCl_2_, 10 mM HEPES, 0.4 mM ethylene glycol-bis-(aminoethyl ethane)-N,N,N’,N’-tetraacetic acid (EGTA), 2 mM MgATP, 0.3 mM GTP-Tris, 10 phosphocreatine, 50mM QX-314 (pH 7.2). For the recordings of mEPSCs, bath solution was supplied with 1 μM tetrodotoxin (TTX, Abcam). mEPSCs from the last 100 s intervals at the end of 5 min recordings with TTX were collected and analyzed using a template based event detection feature of Clampfit 10.2 software. Template was generated from our own data. All signals were digitized at 20 kHz, filtered at 2 kHz, and analyzed offline with Clampfit software (Molecular Devices). All data were exported to and processed in Microsoft Excel

### Data Availability

Source data, summary data and statistical analysis for Figures 3, 4, 5 and 6 provided in Figure Supplements.

## ACKNOWLEDGEMENTS

This work was supported by funding from the Joel G. Hardman Chair in Pharmacology (RBE) and the National Institutes of Health (DA025143, NS094738 and OD00876101 to QZ). 3D-SIM and confocal analyses were performed using the VUMC Cell Imaging Shared Resource (supported by NIH grants CA68485, DK20593, DK58404, DK59637 and EY08126) and training from Dr. Ana M. Carneiro. High-throughput sequence analyses were performed by VANTAGE (Vanderbilt Technologies for Advanced Genomics) and supported by the Vanderbilt Ingram Cancer Center (P30 CA68485), the Vanderbilt Vision Center (P30 EY08126), and NIH/NCRR (G20 RR030956). We also thank Vanderbilt investigators for providing tissues from different species for molecular analysis [Dr. Randy Blakely (*C. elegans*), Dr. Bih-Hwa Shieh (*Drosophila*), Dr. James Patton (zebrafish), and Dr. Christine Konradi (rat)].

### COMPETING INTERESTS

The authors report no competing interest.

## Pearson’s Correlation Coefficients (PCCs) from neurons transfected with edited or non-edited CAPS1-GFP

**Pearson's Correlation Coefficients (PCCs) from neurons transfected with edited or non-edited CAPS1-GFP.**
|  | Edited CAPS1-GFP |  |  | Non-edited CAPS1-GFP |  |  |
| --- | --- | --- | --- | --- | --- | --- |
|  | GFP/PSD | GFP/Syp | PSD/Syp | GFP/PSD | GFP/Syp | PSD/Syp |
| Total number of values | 106 | 106 | 106 | 138 | 138 | 138 |
| Minimum | -0.22 | -0.20 | -0.18 | -0.22 | -0.090 | -0.34 |
| 25% Percentile | -0.006 | 0.40 | 0.013 | 0.033 | 0.34 | 0.049 |
| Median | 0.10 | 0.52 | 0.10 | 0.12 | 0.47 | 0.14 |
| 75% Percentile | 0.17 | 0.64 | 0.22 | 0.23 | 0.57 | 0.21 |
| Maximum | 0.52 | 0.80 | 0.50 | 0.42 | 0.80 | 0.43 |
| Mean | 0.10 | 0.50 | 0.11 | 0.12 | 0.45 | 0.13 |
| Std. Deviation | 0.16 | 0.20 | 0.16 | 0.14 | 0.19 | 0.13 |
| Std. Error of Mean | 0.015 | 0.019 | 0.015 | 0.012 | 0.016 | 0.011 |
| Lower 95% CI of mean | 0.07 | 0.46 | 0.08 | 0.10 | 0.42 | 0.11 |
| Upper 95% CI of mean | 0.13 | 0.54 | 0.14 | 0.15 | 0.48 | 0.15 |

**ANOVA (Mean PCC)**
| *Source of Variation* | *SS* | *df* | *MS* | *F* | *P-value* |
| --- | --- | --- | --- | --- | --- |
| Between Groups | 20.59 | 5 | 4.12 | 155.43 | 3.617E-112 |
| Within Groups | 19.23 | 726 | 0.03 |  |  |
| Total | 39.82 | 731 |  |  |  |

**t-Test: Two-Sample Assuming Equal Variances**
|  | *NE GFP/Syp* | *E GFP/Syp* |
| --- | --- | --- |
| Mean | 0.45 | 0.50 |
| Variance | 0.036 | 0.039 |
| n | 138 | 106 |
| P(T<=t) two-tail | 0.052 |  |

## Statistical analysis of Syp staining (Top) or GFP Staining (Bottom) foci volume presented in Figure 3D

**Statistical analysis of Syp staining (Top) or GFP Staining (Bottom) foci volume presented in Figure 3D.**
| t-Test: Two-Sample Assuming Equal Variances |  |  | F-Test Two-Sample for Variances |  |  |
| --- | --- | --- | --- | --- | --- |
|  | *Non-edited Syp* | *Edited Syp* |  | *Non-edited Syp* | *Edited Syp* |
| Mean | 0.11 | 0.11 | Mean | 0.11 | 0.11 |
| Variance | 0.010 | 0.0086 | Variance | 0.010 | 0.0086 |
| n | 208 | 202 | n | 208 | 202 |
| P(T<=t) two-tail | 0.85 |  | P(F<=f) one-tail | 0.10 |  |
|  | *Non-edited GFP* | *Edited GFP* |  | *Non-edited GFP* | *Edited GFP* |
| Mean | 0.16 | 0.15 | Mean | 0.16 | 0.15 |
| Variance | 0.017 | 0.015 | Variance | 0.017 | 0.015 |
| n | 208 | 202 | n | 208 | 202 |
| P(T<=t) two-tail | 0.16 |  | P(F<=f) one-tail | 0.22 |  |

## Statistical analysis of Syp staining (Top) or GFP Staining (Bottom) foci Area presented in Figure 3E

**Statistical analysis of Syp staining (Top) or GFP Staining (Bottom) foci Area presented in Figure 3E.**
| t-Test: Two-Sample Assuming Equal Variances |  |  | F-Test Two-Sample for Variances |  |  |
| --- | --- | --- | --- | --- | --- |
|  | *Non-edited Syp* | *Edited Syp* |  | *Non-edited Syp* | *Edited Syp* |
| Mean | 1.29 | 1.24 | Mean | 1.29 | 1.24 |
| Variance | 0.83 | 0.63 | Variance | 0.83 | 0.63 |
| n | 208 | 202 | n | 208 | 202 |
| P(T<=t) two-tail | 0.52 |  | P(F<=f) one-tail | 0.026 |  |
|  | *Non-edited GFP* | *Edited GFP* |  | *Non-edited GFP* | *Edited GFP* |
| Mean | 1.67 | 1.46 | Mean | 1.67 | 1.46 |
| Variance | 0.95 | 0.87 | Variance | 0.95 | 0.87 |
| n | 208 | 202 | n | 208 | 202 |
| P(T<=t) two-tail | 0.024 |  | P(F<=f) one-tail | 0.26 |  |

## Statistical analysis of Syp staining (Top) or GFP Staining (Bottom) foci sphericity presented in Figure 3F

**Statistical analysis of Syp staining (Top) or GFP Staining (Bottom) foci sphericity presented in Figure 3F.**
| t-Test: Two-Sample Assuming Equal Variances |  |  | F-Test Two-Sample for Variances |  |  |
| --- | --- | --- | --- | --- | --- |
|  | *Non-edited Syp* | *Edited Syp* |  | *Non-edited Syp* | *Edited Syp* |
| Mean | 0.81 | 0.84 | Mean | 0.81 | 0.84 |
| Variance | 0.0085 | 0.0072 | Variance | 0.0085 | 0.0072 |
| n | 208 | 202 | n | 208 | 202 |
| P(T<=t) two-tail | 0.00079 |  | P(F<=f) one-tail | 0.10 |  |
|  | *Non-edited GFP* | *Edited GFP* |  | *Non-edited GFP* | *Edited GFP* |
| Mean | 0.83 | 0.87 | Mean | 0.83 | 0.87 |
| Variance | 0.0056 | 0.0049 | Variance | 0.0056 | 0.0049 |
| n | 208 | 202 | n | 208 | 202 |
| P(T<=t) two-tail | 2.52E-07 |  | P(F<=f) one-tail | 0.18 |  |

## Statistical analysis of data in Figure 4

One-way ANOVA followed by unpaired t-test was applied for both FM fluorescence intensity and bouton size (Figure 4B). ANOVA on fluorescence, *p* values = 0.32 (NT), 0.18 (E), 0.27 (GFP), 0.011 (NE). Subsequent unpaired t-test between NE and E yielded *p* = 0.008. For ANOVA on bouton size, *p* = 0.34 (NT), 0.30 (E), 0.25 (GFP), 0.029 (NE). Subsequent unpaired t-test between NE and E yielded *p* = 0.04. Decay rate in Figure 4C was analyzed by one-way ANOVA followed by paired t-test. For ANOVA, *p* = 0.30 (NT), 0.06 (E), 0.19 (GFP), and 0.021 (NE). Subsequent unpaired t-test between NE and E yielded *p* = 0.009.

## Summary statistics for data presented in Figure 5C-E

|  | Average |  |  | SE |  |  | F-test |  |  | t-test |  |  |
| --- | --- | --- | --- | --- | --- | --- | --- | --- | --- | --- | --- | --- |
| Figure | NE | E | cntrl | NE | E | cntrl | NE to E | E to Cntrl | NE to cntrl | NE to E | E to Cntrl | NE to cntrl |
| 5C | 0.55 | 0.48 | 0.52 | 0.011 | 0.0081 | 0.0053 | 0.0030 | 0.0054 | 1.56E-09 | 3.33E-06 | 8.16E-05 | 0.039 |
| 5D | 0.18 | 0.19 | 0.17 | 0.0063 | 0.0054 | 0.0037 | 0.10 | 3.17E-04 | 0.063 | 0.27 | 0.49 | 0.023 |
| 5E | 0.28 | 0.33 | 0.31 | 0.0099 | 0.0090 | 0.0052 | 0.44 | 2.50E-09 | 5.98E-08 | 6.79E-05 | 0.011 | 0.027 |

Two tailed t-tests, assuming unequal variance were performed, as determined by F-test, in Microsoft Excel. NE = non-edited, E = edited, cntrl = control.

## Summary data and statistical analysis of data presented in Figure 5F

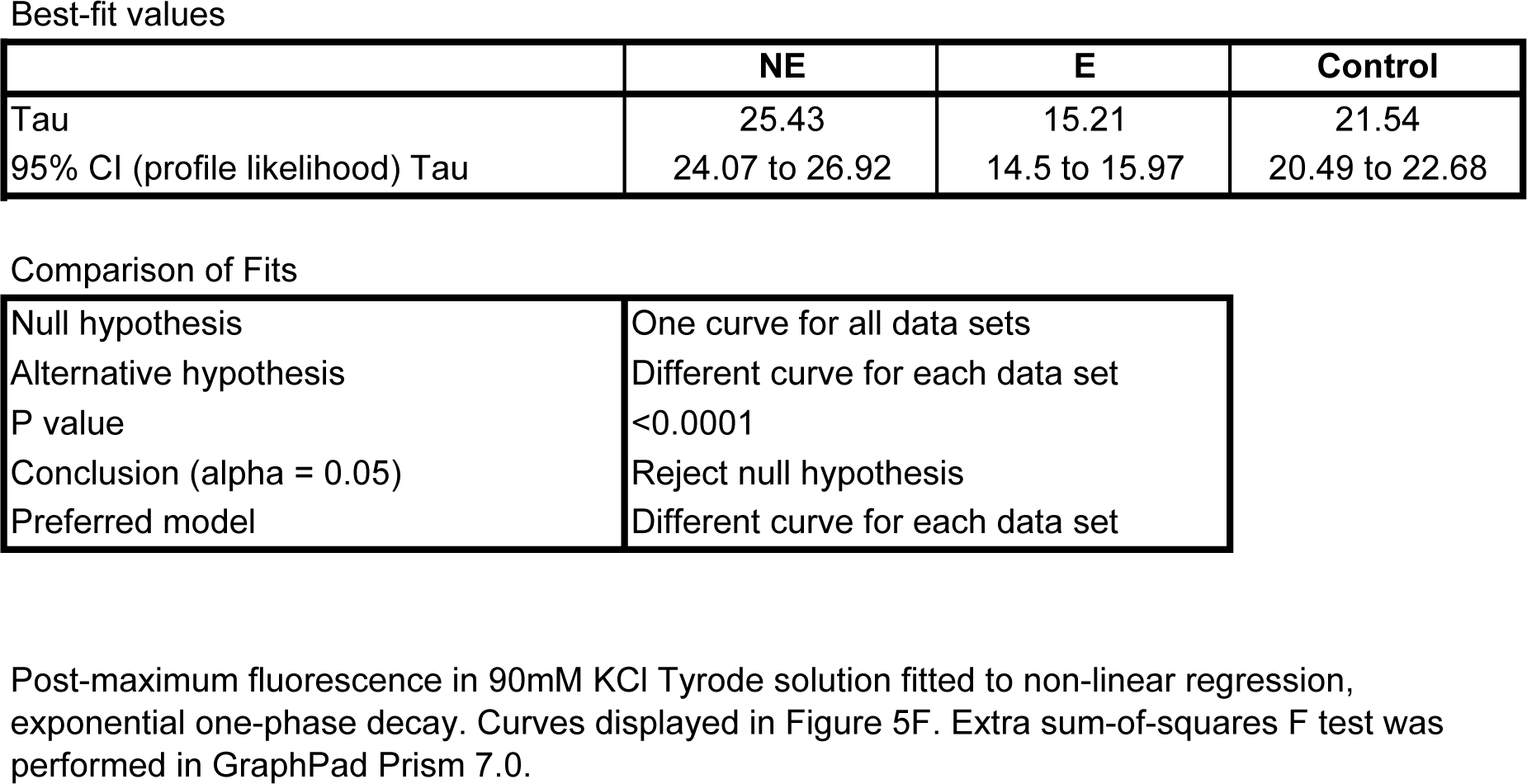

Best-fit values
|  | NE | E | Control |
| --- | --- | --- | --- |
| Tau | 25.43 | 15.21 | 21.54 |
| 95% CI (profile likelihood) Tau | 24.07 to 26.92 | 14.5 to 15.97 | 20.49 to 22.68 |

Comparison of Fits
|  |  |
| --- | --- |
| Null hypothesis | One curve for all data sets |
| Alternative hypothesis | Different curve for each data set |
| P value | <0.0001 |
| Conclusion (alpha = 0.05) | Reject null hypothesis |
| Preferred model | Different curve for each data set |

## Statistical analysis of data presented in Figure 6

**Statistical analysis of data presented in Figure 6.**
| **Figure** | **Test** | **Comparison** | ***p*** |
| --- | --- | --- | --- |
| 6B | K-S | E and NE | 0.022 |
| 6E | Unpaired t-test | E and NE – 0 FRR events | 0.11 |
|  |  | E and NE – 1 FRR events | 0.017 |
|  |  | E and NE – 2 FRR events | 0.024 |
|  |  | E and NE – 3 FRR events | 0.022 |
|  |  | E and NE – 4 FRR events | 0.33 |
|  |  | E and NE – 5 FRR events | 0.40 |
| 6F | Unpaired t-test | E and NE decay rate | 0.35 |
| 6G | K-S | E and NE | 0.00119 |

